# Functional traits fail to predict long-term responses to climate change

**DOI:** 10.1101/2023.07.27.550868

**Authors:** Jeremy D. Ash, Daijiang Li, Sarah E. Johnson, David A. Rogers, Donald M. Waller

## Abstract

**Aim:** Environmental conditions strongly affect the distribution and abundance of species via complex forces. Shifts in environmental conditions and differences in the speed and scale of these effects complicate our efforts to infer how species will respond to future environmental change. We test how 18 functional traits affect plant species responses to gradients in environmental conditions and 50-year shifts in climate.

**Location:** We analyzed 50-year shifts in the distribution and abundance of 153 plant species distributed across 284 sites in Wisconsin, USA.

**Time period:** 1950s to 2000s.

**Major taxa studied:** Vascular plants (much of the flora of NE North America).

**Methods:** We used random forest and integrated hierarchical mixed models to test how plant abundances (and 50-year changes in abundance) track gradients in overstory, soil, and climatic conditions.

**Results:** Within study periods, plant abundances reflect gradients in environmental conditions. Leaf traits affected local abundance (both directly and via trait-environment interactions) in the 1950s and 2000s. Strong soil and temperature effects in the 1950s have weakened while precipitation effects have strengthened. Although we expected these models to also predict how plants would respond to shifts in climate, they did not.

**Main conclusions:** Lags in species’ responses, increases in the stochastic forces affecting community assembly, and other forces limit the ability of models fitted to static data (e.g., space-for-time substitutions) to predict how plant species will respond to long-term shifts in environmental conditions. We must therefore be cautious about applying trait-based species distribution models to predict how climate change will affect species distributions and community structure.

## Introduction

Population and community ecologists seek to characterize the factors determining variation in species abundance and distribution. These factors include how species’ characteristics (functional traits) affect their response to gradients in environmental conditions, e.g., species distribution models (Franklin, 2010). Such models help us understand current species’ distributions as well as predict future shifts in distribution (Elith & Leathwick, 2009). With global changes in climate, many species are shifting their distribution toward the poles or higher in elevation (Parmesan & Yohe, 2003). Species, however, vary greatly in how they respond to shifts in environmental conditions (Ibáñez *et al*., 2013). Inferring these effects of climate change is complicated by the effects of simultaneous changes in atmospheric CO2 levels (Cole et al. 2010), aerial nitrogen deposition (Clark et al. 2019; Staude et al. 2020), disturbance regimes (Nowacki & Abrams, 2014; Stevens *et al*., 2015), and other environmental factors (Austin & Van Niel, 2011).

Do differences in response among species reflect plant functional traits? Species’ traits illuminate the mechanisms underlying how species respond to drivers of global change. If species that share traits respond similarly we could use traits to predict their responses (Buckley & Kingsolver, 2012; Brown *et al*., 2016). Exploring how, and how consistently, traits affect species’ responses to in environmental conditions could thus help us predict species’ responses to the different kinds of environmental change (Pollock *et al*., 2012). In a global analysis of 17 traits for >20,000 species, Joswig et al. (2022) found trait variation to parallel variation in both climatic and soil factors.

Trait-based approaches are now routinely used to study how environmental conditions affect community assembly (Cornwell & Ackerly, 2009; Laughlin *et al*., 2011; Frenette-Dussault *et al*., 2012; Rolhauser et al. 2021, Beck et al. 2022). Most such studies, however, rely on one-time ‘snapshot’ data or focus on local dynamics. Two exceptions explore trait-environment relationships to understand how these affect long-term shifts in plant species’ abundances. Li et al. (2015) found that traits strongly predict shifts in subtropical tree communities while Soudzilovskaia *et al*. (2013) found similar predictive power in alpine meadows. Many traits strongly affect species responses at broad spatial scales across environmental gradients (Wright *et al*., 2005a; 2005b; Moles *et al*., 2014). How these relationships affect shifts in local and regional plant communities is less clear. Because trait-based community assembly can depend on spatial scale (Cavender-Bares *et al*., 2006; Swenson & Enquist, 2009; Kraft & Ackerly, 2010; Xing *et al*., 2014), our efforts to link traits to species’ responses should explicitly consider scale and spatial extent.

Trait-based community ecologists often use "fourth corner" approaches to understand how species’ traits respond to environmental gradients (McCune *et al*., 2002; Dray & Legendre, 2008; Dray *et al*., 2014). This approach employs three data matrices: species by site (from either species’ incidence or abundance); site by environmental variables; and species by trait values. These relationships are then combined to infer a fourth “missing corner” matrix to indicate how traits relate to environmental conditions. This approach yields useful relational and descriptive tools but ignores two important questions: 1. How do environmental variables mechanistically affect species’ distributions? 2. Do static associations between traits and environmental variables serve to predict temporal species’ dynamics (Jamil *et al*., 2013)? Jamil et al. (2012) and Pollock et al. (2012) applied hierarchical models to estimate how traits and trait-environmental interactions act to structure species’ distributions allowing them to make stronger inferences than those generated from classical species’ distribution models.

Here we ask how the distributions of forest understory plant species in Wisconsin are structured by environmental gradients and to what extent traits serve to predict species-environment relationships. Despite dramatic shifts in taxonomic diversity (Rooney et al. 2004; Rogers et al. 2008; Johnson et al. 2014; Li and Waller 2015) and strong trait-environment relationships (Amatangelo *et al*., 2014), Wisconsin forest plant communities have experienced little aggregate change in functional diversity (Sonnier *et al*., 2014). To assess the consistency of species-environment relationships and how well traits predict responses to environmental variation, we combined baseline data derived from extensive surveys by J.T. Curtis and his students in the 1950s (Curtis, 1959) with re-surveys of the same sites in the 2000s (Waller *et al*., 2012). We use these data to construct separate species’ distribution models for each survey period. We then construct a third model to predict changes in the abundance of each species between the two periods. These three models allow us to assess: 1) how site-level changes in environmental conditions affect spatial patterns of species’ abundance; 2) whether the dominant traits and environmental gradients structuring species’ distributions have remained stable over the past 50 years; and 3) whether these trait-environment relationships allow us to predict how species are responding to shifts in environmental conditions. We are fortunate here to be able to exploit and combine exceptionally rich sets of data on species distribution and abundance, life history variation, functional traits, and local environmental conditions. These manifold data lend statistical strength to our analyses and allow us to explore whether functionally similar species respond similarly to environmental gradients and change.

We focus here on three kinds of abiotic data: soil conditions, successional status, and climatic gradients. Variation in soil chemistry and texture is a key factor structuring plant communities (Hutchinson *et al*., 1999; Small & McCarthy, 2005; Gilliam, 2014). Disturbance-driven succession also strongly affects community composition in Wisconsin forests (Rooney *et al*., 2004; Wiegmann & Waller, 2006; Rogers *et al*., 2008; Johnson *et al*., 2014; Li & Waller, 2015). Considerable changes in temperature and precipitation have also occurred over the past half century in Wisconsin including substantial spatial variation in these trends (Kucharik *et al*., 2010; WICCI, 2011). If the forces affecting species distributions have changed over the past 50 years (or the rates and extents of species’ responses to these forces), species’ distributions might well show complex patterns of response to a changing set of environmental variables.

We previously described broad geographic shifts in species’ distribution that largely parallel changes in climate (Ash *et al.*, 2017). These species also show fairly consistent trait-environment relationships even as their communities have shifted substantially in composition and relative abundance (Amatangelo *et al*., 2014). The results we present here complement these studies by analyzing site-level patterns in more detail (using stronger statistical methods) to illuminate how traits affect shifts in species abundance in response to changing local environmental conditions. As noted above, the set of traits driving species-environment relationships may have changed. This seems especially likely given that the factors driving ecological change differ considerably among the four community types we analyze (Rooney *et al*., 2004; Rogers *et al*., 2008; 2009; Johnson *et al*., 2014; Li & Waller, 2015). Stochastic neutral processes reflecting local abundance and dispersal (Hubbell 2001) also appear to have increased in importance relative to deterministic trait-based niche processes (Waller et al. 2017).

## Methods

### Study system

To encompass a broad range of geographic and environmental variation, we analyzed changes across 284 sites distributed among four community types in Wisconsin (Fig. 1): pine barrens of the central sand plains (CSP, n = 30), northern upland forests (NUF, n= 96), southern lowland forests (SLF, n = 40) and southern upland forests (SUF, n = 118). Various ecological changes have affected these communities. Species susceptible to deer herbivory have declined conspicuously in most NUF communities while a few exotic species and species that avoid or tolerate deer herbivory (e.g., grasses and sedges) have greatly increased (Rooney *et al*., 2004; Wiegmann & Waller, 2006). Exotic species invasions and declines in native species are more dramatic in the more fragmented upland forests of southern Wisconsin (Rogers *et al*., 2009). Fire suppression, succession, and consequent “mesification” also affect many SUF sites, but these processes are especially conspicuous in the Pine barren (CSP) communities (Li & Waller, 2015). Southern lowland forest communities have become more diverse yet more similar reflecting increases in habitat connectivity and highly regulated hydrological regimes (Johnson *et al*., 2014).

**Figure 1.**
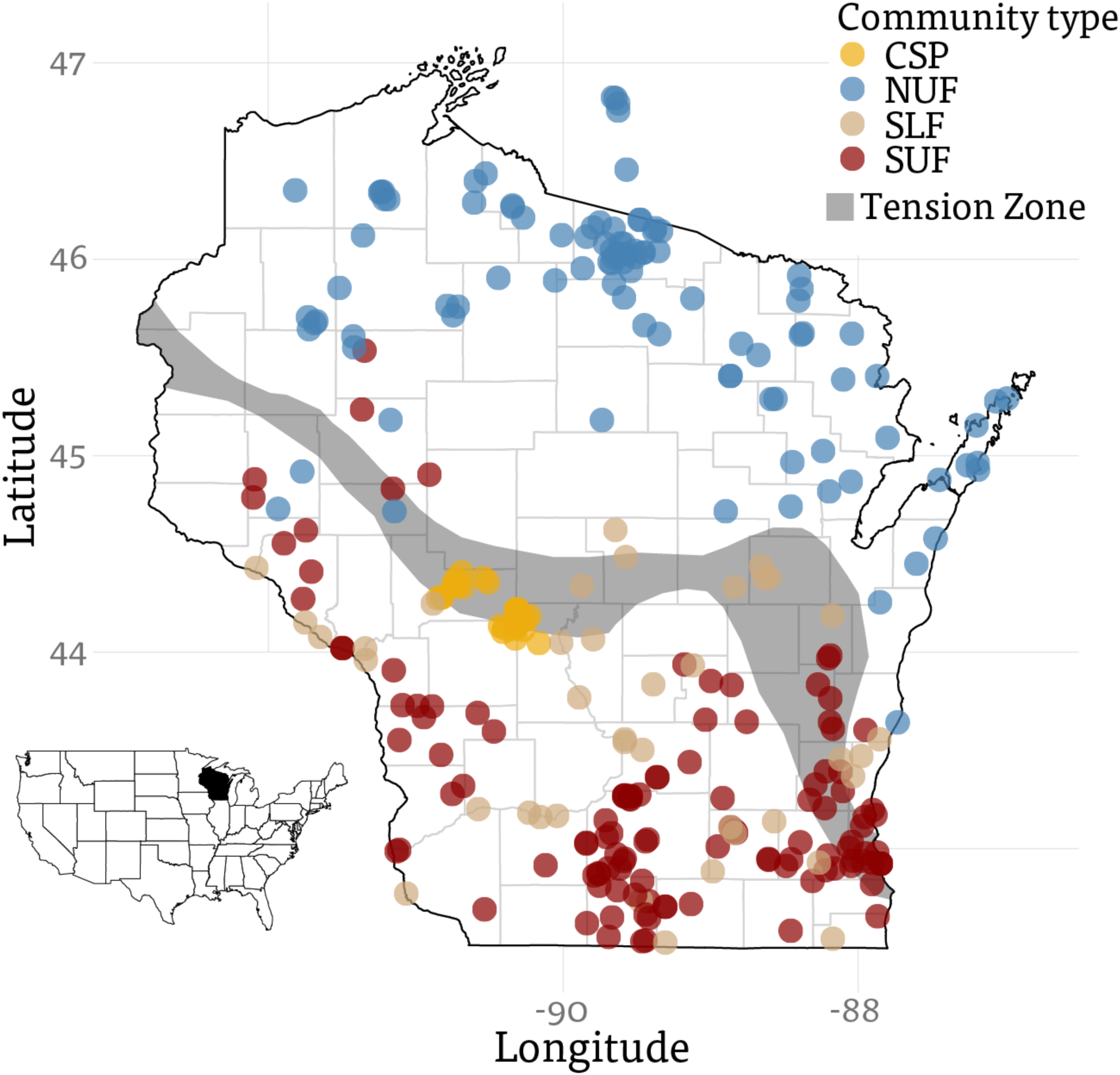
Map showing locations of the surveyed sites. Community types are color coded: CSP: pine barrens of central sand plains; NUF: northern upland forests; SLF: southern lowland forests; SUF: southern upland forests.

We briefly describe the original sampling and resurvey methods here. More detailed descriptions appear in the publications cited above. J.T. Curtis and colleagues first surveyed >1000 plant communities across Wisconsin in the 1950s (Curtis, 1959). They chose sites exhibiting minimal disturbance (i.e., logging) and fragmentation (i.e., sites >6 ha). Curtis-era samples relied on 20 (rarely 40) evenly-spaced 1 m^2^ quadrats generally arranged along a square or U-shaped transect. Resurveys began in the 2000s applying similar but more intensive samples (42-120 1 m^2^ quadrats) to the same understory communities (Waller *et al*., 2012; http://www.botany.wisc.edu/PEL/). The original sites were not permanently marked but re-located accurately using detailed site descriptions and maps (Waller *et al*., 2012). Such resurveys reflect “semi-permanent” plots in the lexicon of ForestReplot (https://forestreplot.ugent.be/).

Within each quadrat, researchers recorded the presence of all vascular plant species (herbs, shrubs and trees) allowing estimates of abundance as the frequency (sum of occupied quadrats) of each species at each site. Here, we focus on species occurring in 10+ sites in one or both survey periods and for which we have full coverage of functional traits (see below). These filters yielded a residuum of 153 species.

### Environmental Variables

We used 8 km gridded climate data spatially interpolated from the extensive network of weather stations covering Wisconsin to characterize shifts in climate between 1950 and 2006 (Kucharik *et al*., 2010). We computed 19 biologically meaningful BIOCLIM variables from daily precipitation and minimum and maximum temperatures from each of our 284 sites based on five-year means for both survey periods (1950-1954 and 2000-2004 – Table 1). Using 5-year means smooth inter-annual variability and partially accounts for potential lags in how species respond to climate change (DeFrenne *et al*., 2013). These summarize biologically relevant climate variables and seasonal and annual aspects of climate change (Booth *et al*., 2013). They include mean temperature and precipitation over each season and energy-based measures like potential evapotranspiration (PET) calculated using the Thornthwaite equation (Thornthwaite, 1948). We estimated annual water deficit as the difference between potential evapotranspiration and annual precipitation. We inferred day-length from site latitude (Forsythe *et al*., 1995) and calculated growing degree days using 10°C as the base threshold. We used the *dismo* package in R to calculate the 19 summary variables (Hijmans *et al*., 2015).

**Table 1.** The 34 climate and other environmental variables considered in the first stage of modeling and their abbreviations.

| Variable | Explanation | Units |
| --- | --- | --- |
| BIO1 | Annual Mean Temperature | °C |
| BIO2 | Mean Diurnal Range (Mean of monthly (max - min temp)) | °C |
| BIO3 | Isothermality (BIO2/BIO7) (* 100) | unitless |
| BIO4 | Temperature Seasonality (standard deviation *100) | °C |
| BIO5 | Max Temperature of Warmest Month | °C |
| BIO6 | Min Temperature of Coldest Month | °C |
| BIO7 | Temperature Annual Range (BIO5-BIO6) | °C |
| BIO8 | Mean Temperature of Wettest Quarter | °C |
| BIO9 | Mean Temperature of Driest Quarter | °C |
| BIO10 | Mean Temperature of Warmest Quarter | °C |
| BIO11 | Mean Temperature of Coldest Quarter | °C |
| BIO12 | Annual Precipitation | mm |
| BIO13 | Precipitation of Wettest Month | mm |
| BIO14 | Precipitation of Driest Month | mm |
| BIO15 | Precipitation Seasonality (Coefficient of Variation) | unitless |
| BIO16 | Precipitation of Wettest Quarter | mm |
| BIO17 | Precipitation of Driest Quarter | mm |
| BIO18 | Precipitation of Warmest Quarter | mm |
| BIO19 | Precipitation of Coldest Quarter | mm |
| precip_fall | Fall precipitation | mm |
| precip_win | Winter precipitation | mm |
| precip_spr | Spring precipitation | mm |
| precip_sum | Summer precipitation | mm |
| temp_fall | Mean fall temperature | °C |
| temp_win | Mean winter temperature | °C |
| temp_spr | Mean spring temperature | °C |
| temp_sum | Mean summer temperature | °C |
| water_def | Yearly water deficit (PET – BIO12) | mm/yr |
| gdd | Growing degree days (base temperature 10 °C) | °C |
| tmin_win | Mean daily minimum winter temperature | °C |
| pet | Potential evapotranspiration | mm/yr |
| Mean BA | Mean basal area of overstory trees | m<sup>2</sup> |
| Soil (PCA1) | PCA axis 1 score, correlated with pH and cation content | unitless |
| Soil (PCA2) | PCA axis 2 score, correlated with soil texture | unitless |

To characterize soil conditions, we collected and combined 3-10 soil samples from semi-random locations at each site. We then combined these samples and tested for nine soil variables: pH; % organic matter; % sand, clay and silt; and concentrations of phosphorous, potassium, calcium and magnesium (in ppm). We applied Principal Components Analysis to reduce dimensionality of the data focusing on the first two axes that together account for 67% of the total variation in soil conditions among sites. We inverted values on axis 1 so these increase in parallel with soil cations and pH. The second axis largely reflects soil texture, ranging from coarse, sandy soil (low scores) to fine-textured soils (high scores). We only had reliable soil assays from the 2000s survey. We therefore used these values in models in both periods, implicitly assuming that soil texture and chemistry has not changed substantially between the 1950s and 2000s. To incorporate variation in canopy conditions (e.g., successional status), we used the mean basal area (BA) of the trees measured at each site as an indicator of canopy cover. Areal BA estimates would be preferable, but the Curtis-era and 2000s SUF surveys used plotless methods that provide unreliable estimates of basal area and stem density per ha (Waller *et al*., 2012). We re-centered all environmental variables on zero and scaled by standard deviations to facilitate comparisons.

### Functional traits

We measured a broad suite of functional traits to capture a range of life history and ecological strategies (**Table 2**). Trait means for each species reflect measurements on at least 12 individuals (four individuals from each of three sites) following standardized protocols (Pérez-Harguindeguy *et al*., 2013; Waller et al. 2021). These data include estimates of genome size from Wisconsin plants (Bai *et al*., 2012) and each species’ Coefficient of Conservatism (CC), an estimate of habitat fidelity (Swink and Wilhelm 1994; Table 2). Values of CC range from 0 (no fidelity) to 10 (scarce species confined to specific, high-quality habitats). We also included each species’ initial geographic range (a convex hull around sites occupied in the 1950s). All continuous trait variables were centered and scaled by subtracting the mean and dividing by two standard deviations (Gelman 2008). This allows us to easily compare effect sizes among these heterogeneous binary and continuous variables.

**Table 2.**
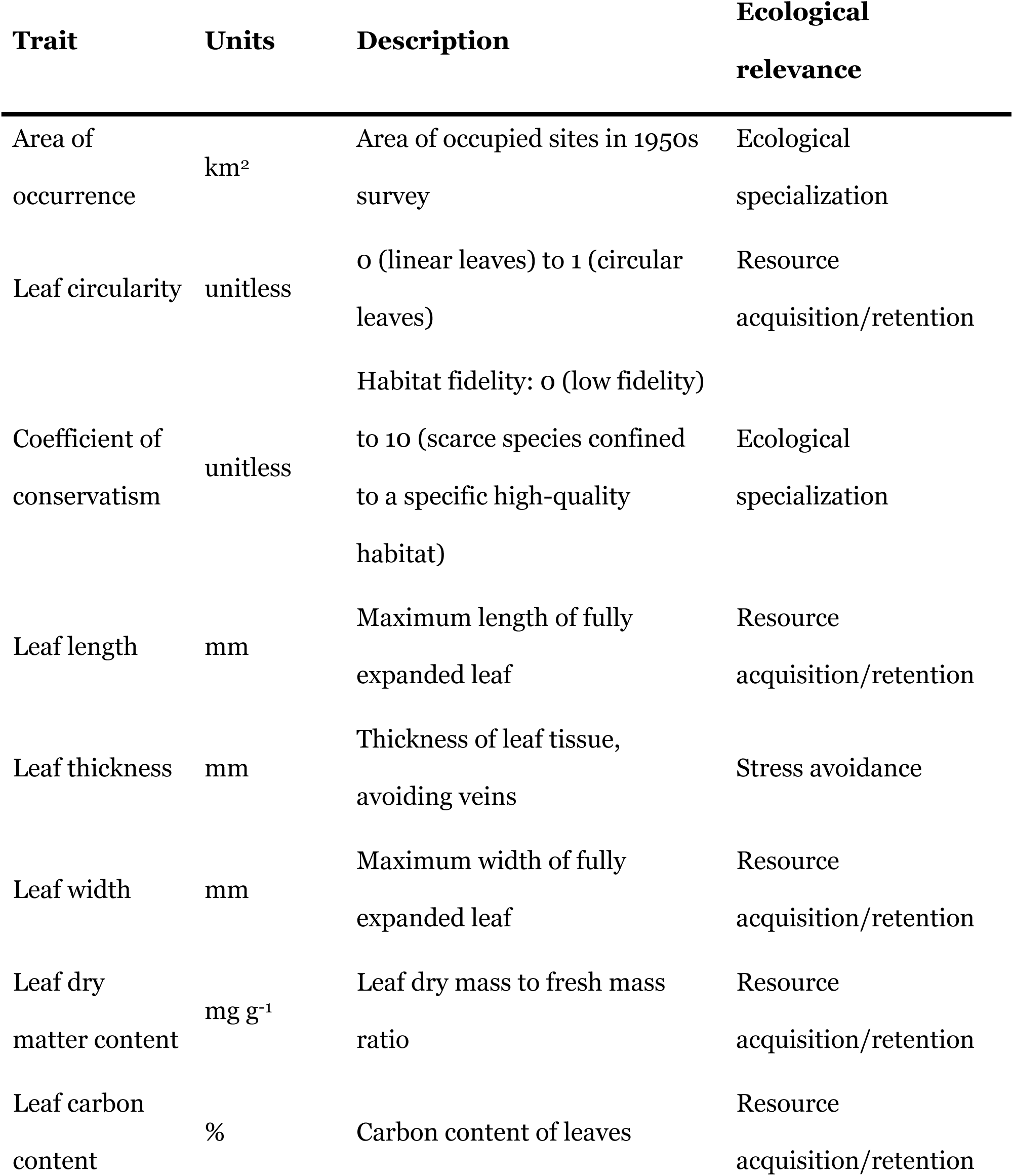
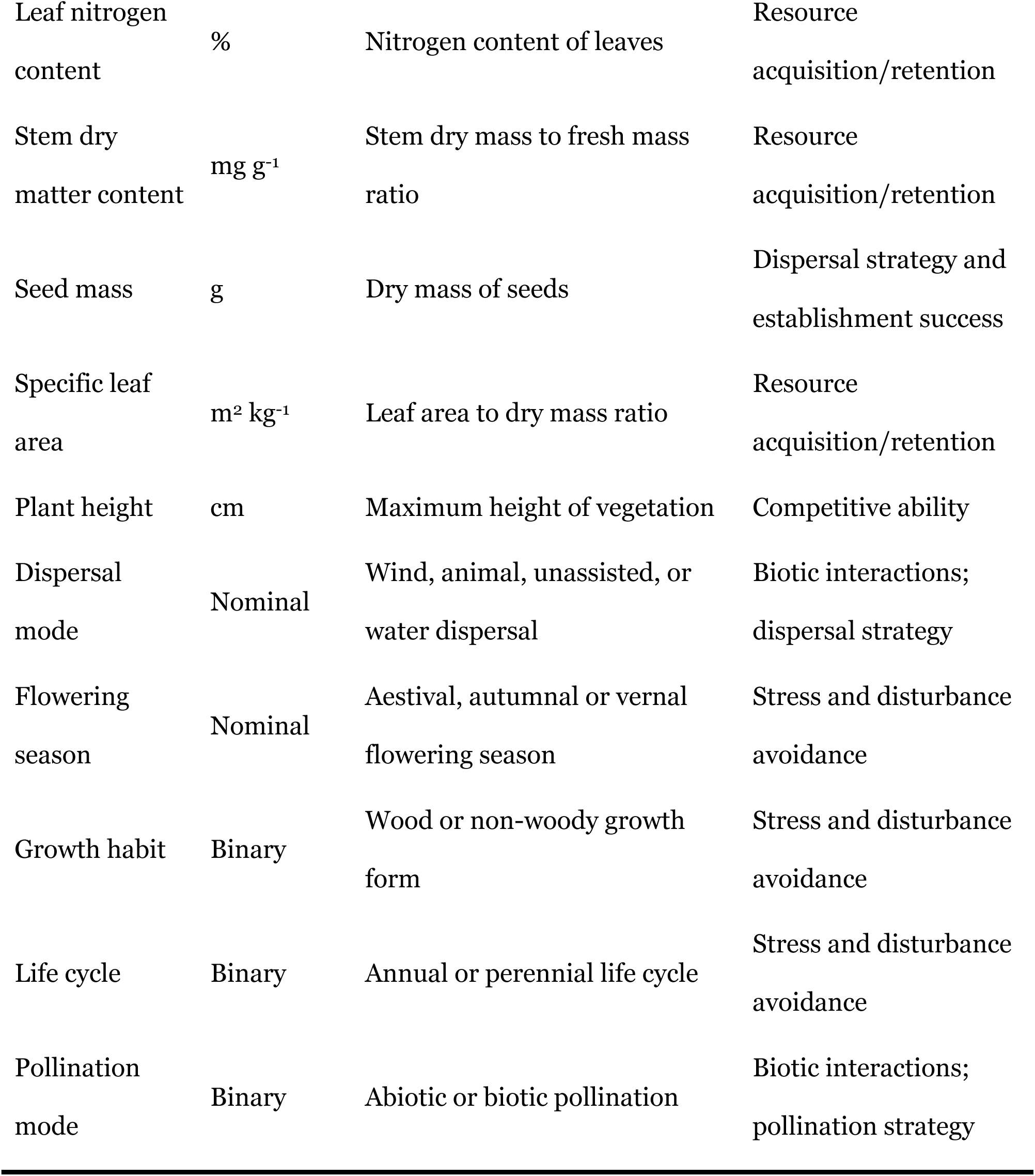
The 18 traits tested in the initial models along with a summary of their ecological meaning or relevance.

### Data Analyses

We analyzed species’ abundances across sites within each survey period and changes in abundance between periods. Within survey periods, we modeled species’ abundances as binomial processes based on the proportion of quadrats occupied, applying a logit transformation: log(y/[1-y]), where y is the proportion of quadrats each species occupied at each site to normalize these responses (Warton & Hui, 2011). To measure changes in abundance for each species at each site, we subtracted these logit-transformed proportions (2000s – 1950s).

To reduce collinearity among the many predictors, we first sought to systematically identify less inter-correlated subsets of variables for each model. For trait-based models of plant abundance in the 1950’s and 2000’s, we applied linear mixed models relating variation in species abundance across sites to each trait variable, including site as a random effect. We then ranked models according to how much variance in species’ abundances they explained, allowing us to select the three traits that best accounted for abundance variation within each species. We applied the same procedure to analyze effects of traits on changes in abundance between the 1950s and 2000s. The explanatory power of these models can be summarized as conditional and marginal R^2^ values referring to all variables or just the fixed factors, respectively (Nakagawa and Schielzeth 2013).

We applied random forest models to analyze how environmental predictor variables affected species abundances in both periods (Table 1). This machine learning method is commonly applied in regression and classification trees (Hastie *et al*., 2009) and to modeling species’ distributions (Prasad *et al*., 2006). We replicated the procedure 500 times to reliably infer which environmental variables most consistently served predict species’ abundances across sites in the two time periods (Figs. S1 – S2). We also analyzed changes in abundance as a function of changes in mean tree basal area and the bioclimatic variables (2000s values minus 1950s values – Fig. S3). As we lacked soil data from the 1950s, we could not analyze effects of changes in soil predictors (Soil PCA1 and 2) in this delta abundance model. This is unfortunate, as these had important effects on plant abundances in the 1950s and 2000s (see Discussion). We then compared species to determine which environmental variables most consistently served to predict species’ distribution patterns. After eliminating collinear variables (correlation > 0.6), we chose the five most important variables for each of the three models. This number reduced overfitting and the number of convergence warnings.

Finally, we applied hierarchical models similar to those recently used to pursue similar questions in other systems (Gelfand *et al*., 2005; Pollock *et al*., 2012; Jamil *et al*., 2013). Our approach matches methods described in Pollock et al. (2012). These use generalized linear mixed effect models (GLMM, Gelman & Hill, 2007) but integrate two steps into one model. First, we fit a regression model for each species, relating their abundance to the environmental variables. Second, we regressed parameters from this first model onto species’ trait values to explore how well traits predict the species-environment relationship. These integrative models include interaction terms to account for the trait-environment relationships while controlling for site as a random effect.

For each survey, we modeled the response of species frequency (the proportion of quadrats occupied, our index of abundance) to the selected environmental variables, species’ traits, and their interactions using a binomial (logit-link) generalized linear mixed model (GLMM). We also included main effects of the four community types (CSP, NUF, SLF, and SUF) and the random effect of site to account for the nested (hierarchical) nature of our data. For the abundance-change model, we modeled the change in logit-transformed proportions as a function of species’ traits, change in the environmental variables, and interactions between these in a linear mixed model assuming a Gaussian error distribution including the same community and site effects. Here, species’ responses to environmental variables are modeled as a random effect, meaning that the average response (intercept) and partial responses to environmental gradients (slopes) can all vary across species. To test the hypothesis that trait-environment relationships declined between the 1950s and 2000s, we applied the 1950s model to the 2000s data and compared coefficients between periods.

We used R (R Development Core Team, 2015) for all analyses, including the following packages: *lme4* for mixed modeling, *performance* for summary *R^2^* statistics from the mixed models (Lüdecke et al. 2021), *randomForest* for random forest modeling (Liaw & Wiener, 2002), *ncdf4*, *raster* and *dismo* for processing climate variables (Pierce, 2014; Hijmans, 2015; Hijmans *et al*., 2015), and *ggplot2* for plotting (Wickham, 2009).

## Results

### Species’ abundance

The random forest models reveal how a few key environmental variables appear to strongly affect variation in species’ abundance across sites (Fig. 2). Soil conditions, in particular, emerged as the most accurate predictors of species’ abundances in both the 1950s and 2000s (Figs. S1 - 2). Growing season temperatures, potential evapotranspiration, and growing degree days appear to have also affected species’ abundances in both survey periods. Although mean tree basal area strongly affected species’ abundances in the 1950s, it became far less important by the 2000s (Figs. 2, S1, and S2). Declines in mean tree BA variance might account for this, but this only occurred at the Central Sand Plains sites (where the S.D. declined from 0.16 to 0.02). Key climate variables also shifted between periods. Measures of temperature important in the 1950s lost importance relative to precipitation variables by the 2000s. Interestingly, changes in mean tree basal area and many of the climate variables affecting abundance in the 2000s appeared to most affect changes in abundance between periods (Fig. S3).

**Figure 2.**
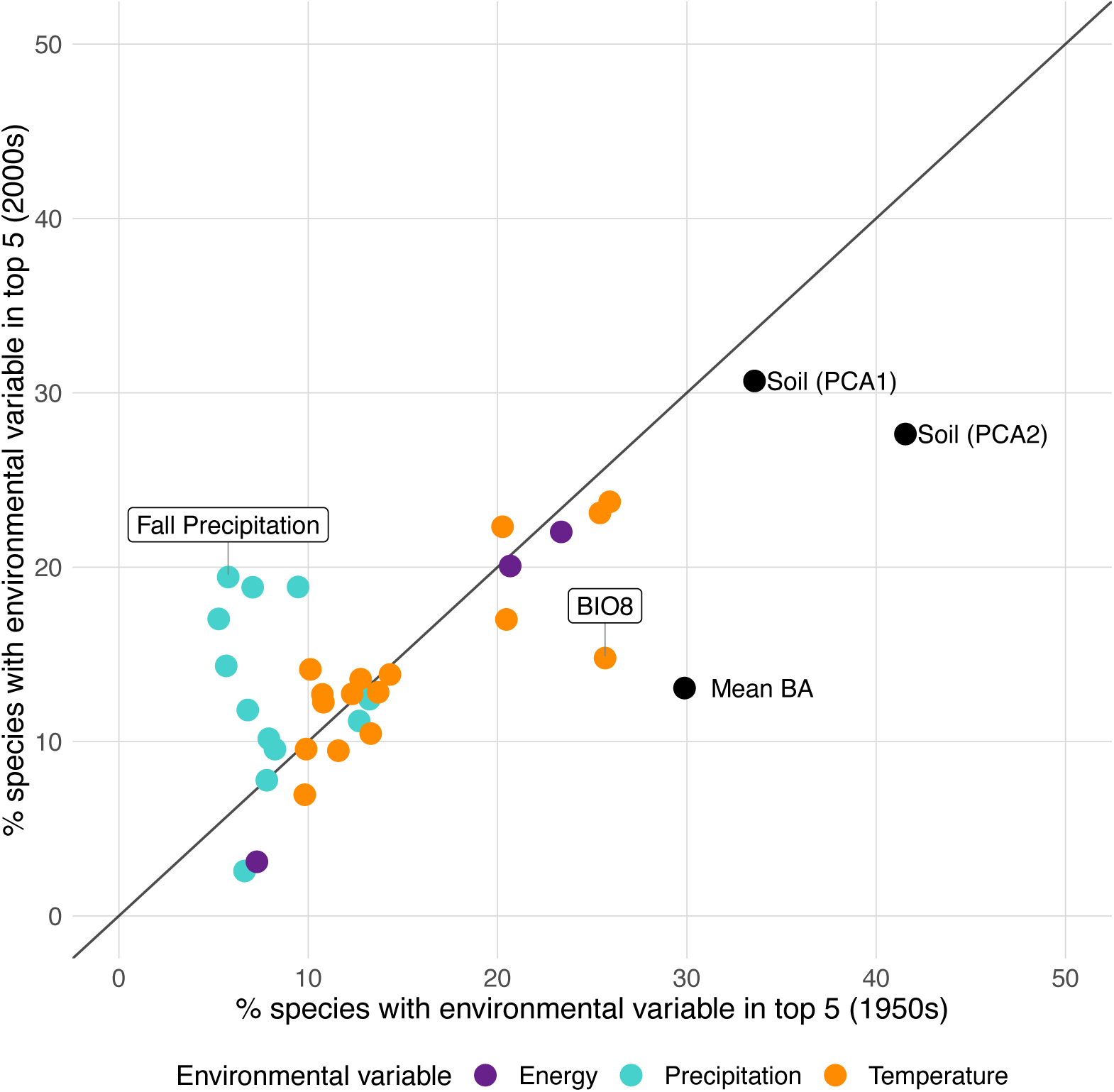
Covariation in the relative importance of environmental variables for predicting species’ abundances in the two time periods (one-to-one line shown for comparison). Values are derived from random forest models within each time period and represent the percent of species where each particular variable was included among the top five predictors explaining species’ distributions. Points are color-coded to distinguish soil variables (PCA axes 1 and 2), and inferred light (mean overstory tree basal area, Mean BA) from climate and energy variables. Climate variables are based on temperature and precipitation alone (separate colors) while the composite energy variables include potential evapotranspiration, growing degree days, and annual water deficit. Note how soil and mean BA have become less important since the 1950s for many species while precipitation variables have grown more important.

Among traits, leaf thickness and dry matter content most consistently affected species’ abundance patterns and how these changed between periods (Tables 3 - 4, Fig. 5). Area of initial occurrence (1950s model), leaf nitrogen content (2000s model), and specific leaf area (change in abundance model) also appeared to affect variation among sites in species’ abundance. These models had strong overall explanatory power with conditional *R^2^* values of 0.61 and 0.70 for 1950s and 2000s abundances, respectively. Marginal *R^2^* values (reflecting fixed effects only) accounted for about a quarter of this (0.16 and 0.17, respectively).

**Table 3.**
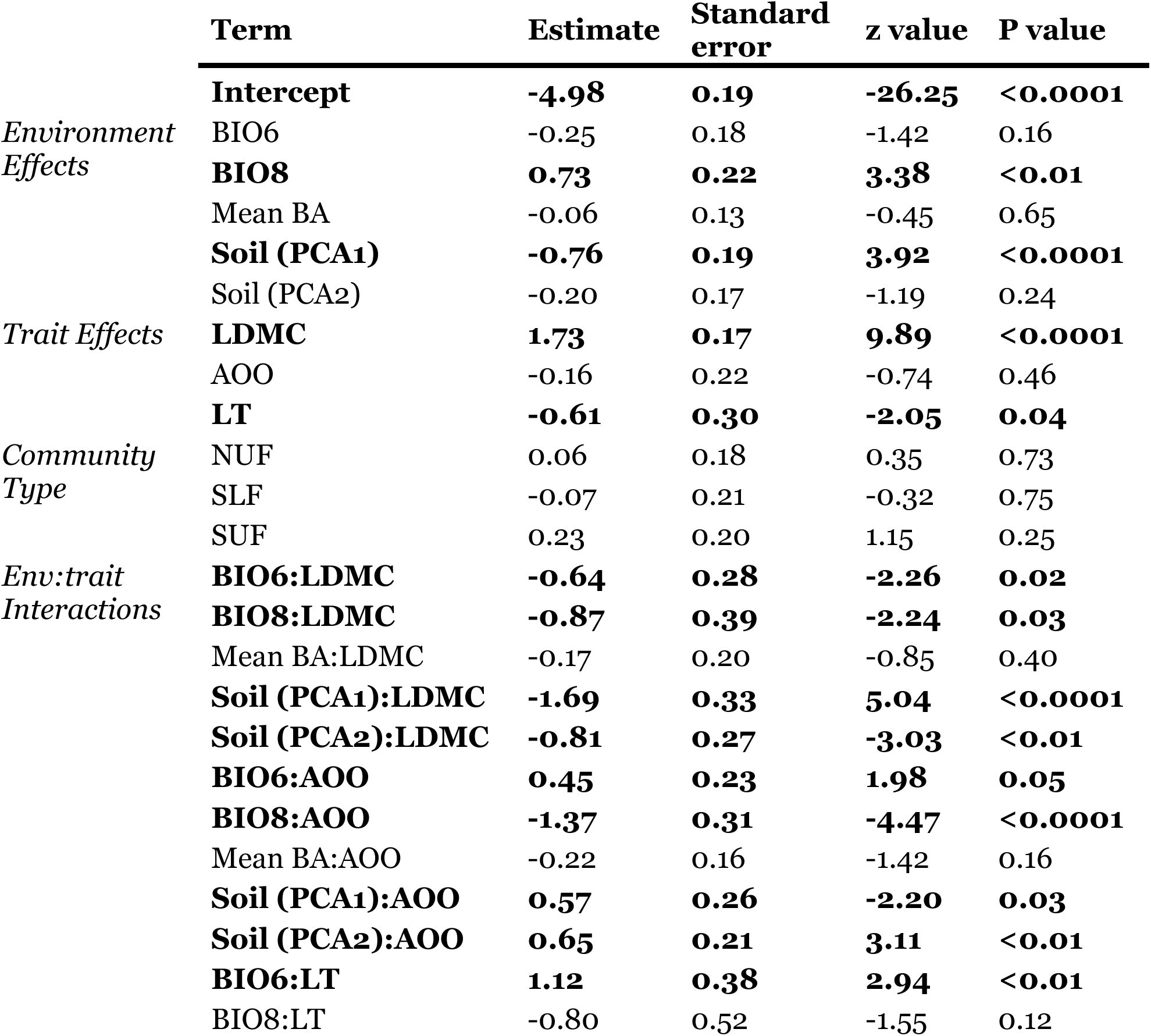
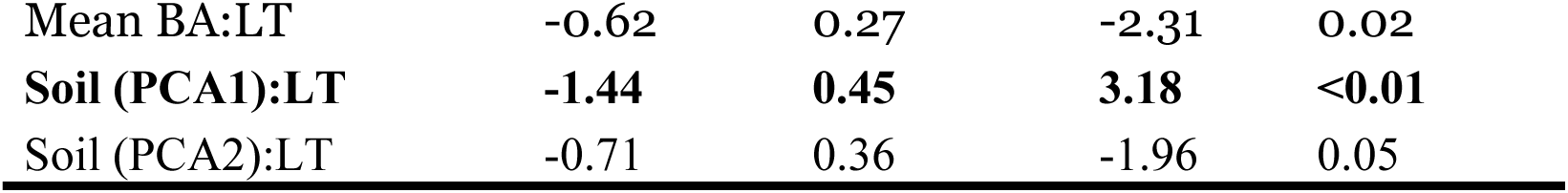
Summary of the **1950s model** showing effects of predictor variables on the abundance (frequency) of 153 species distributed across 284 sites. Model coefficients reflect how each predictor affects estimated abundance on the logit scale. Conditional and marginal R^2^ values are 0.611 and 0.158, respectively, for this model. Abbreviations are: *Environment* - BIO6: minimum temperature of coldest month; BIO8: mean temperature of wettest quarter; Mean BA: mean basal area of overstory trees; and *Traits* - LDMC: leaf dry matter content; AOO: area of occurrence; LT: leaf thickness. Significant effects (p < 0.05) are bolded.

**Table 4.**
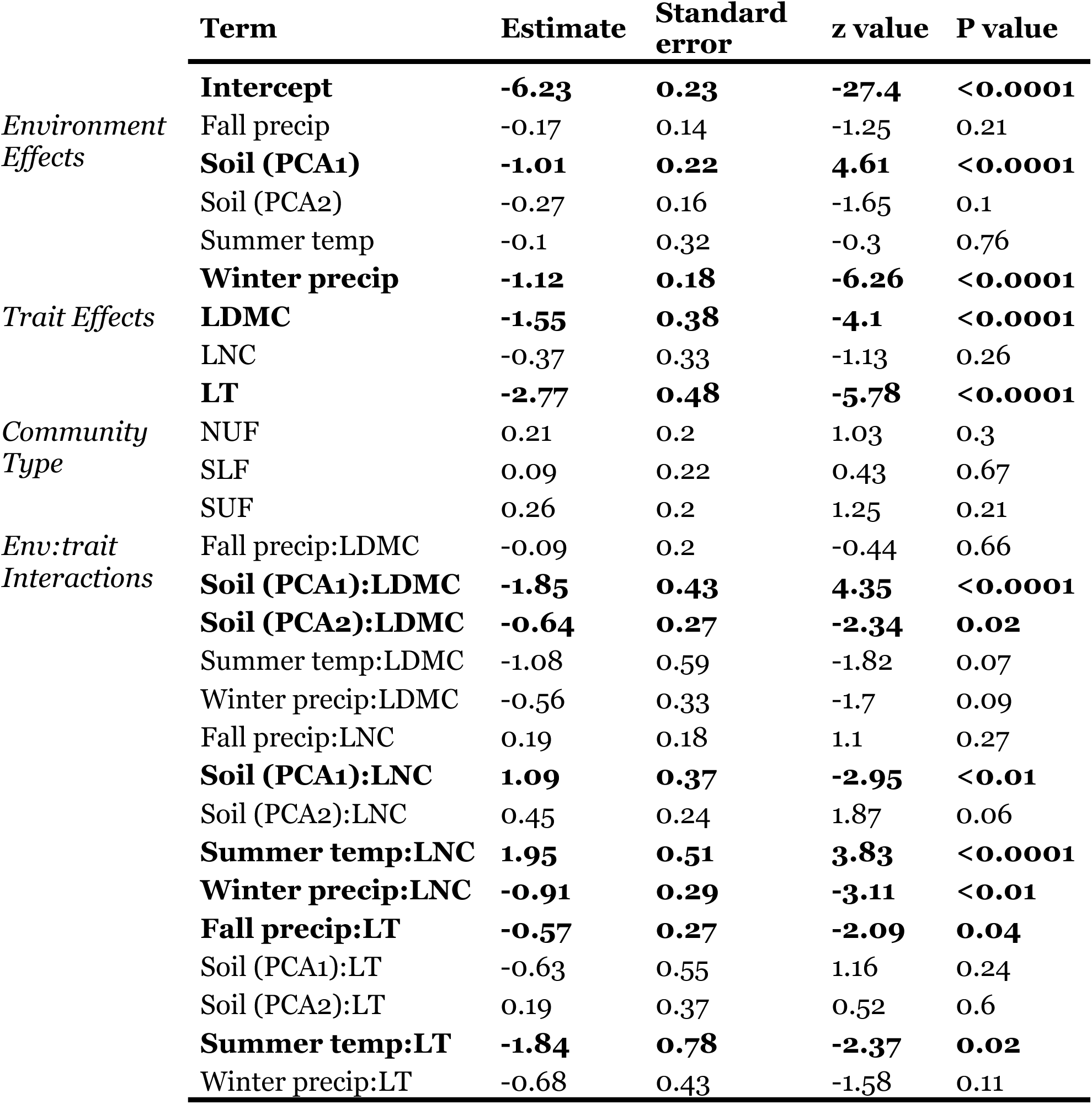
Summary of the **2000s model** showing effects of predictor variables on the abundance (frequency) of 153 species distributed across 284 sites. Model coefficients reflect how each predictor affects estimated abundance on the logit scale. Conditional and marginal R^2^ values are 0.704 and 0.171, respectively, for this model. Trait abbreviations are: LDMC: leaf dry matter content; LNC: leaf nitrogen content; LT: leaf thickness. Significant effects (p < 0.05) are bolded.

### Static models of distribution and abundance

Species vary greatly in how they respond to gradients in environmental conditions, as expected if they are differentially adapted to those conditions (Figs. 3-4). Species individualistic responses vary independently from their abundance. All environmental variables have some power to predict species distribution and abundance, however the relative strength of their effects varies widely across species. Collectively, species clearly respond most strongly to soil chemistry and texture (Soil PCA1 and PCA2) in both time periods, followed by mean temperature of the wettest quarter (BIO8) in the 1950s and mean summer temperature (BIO10) in the 2000s (Figs. S1 and S2).

**Figure 3.**
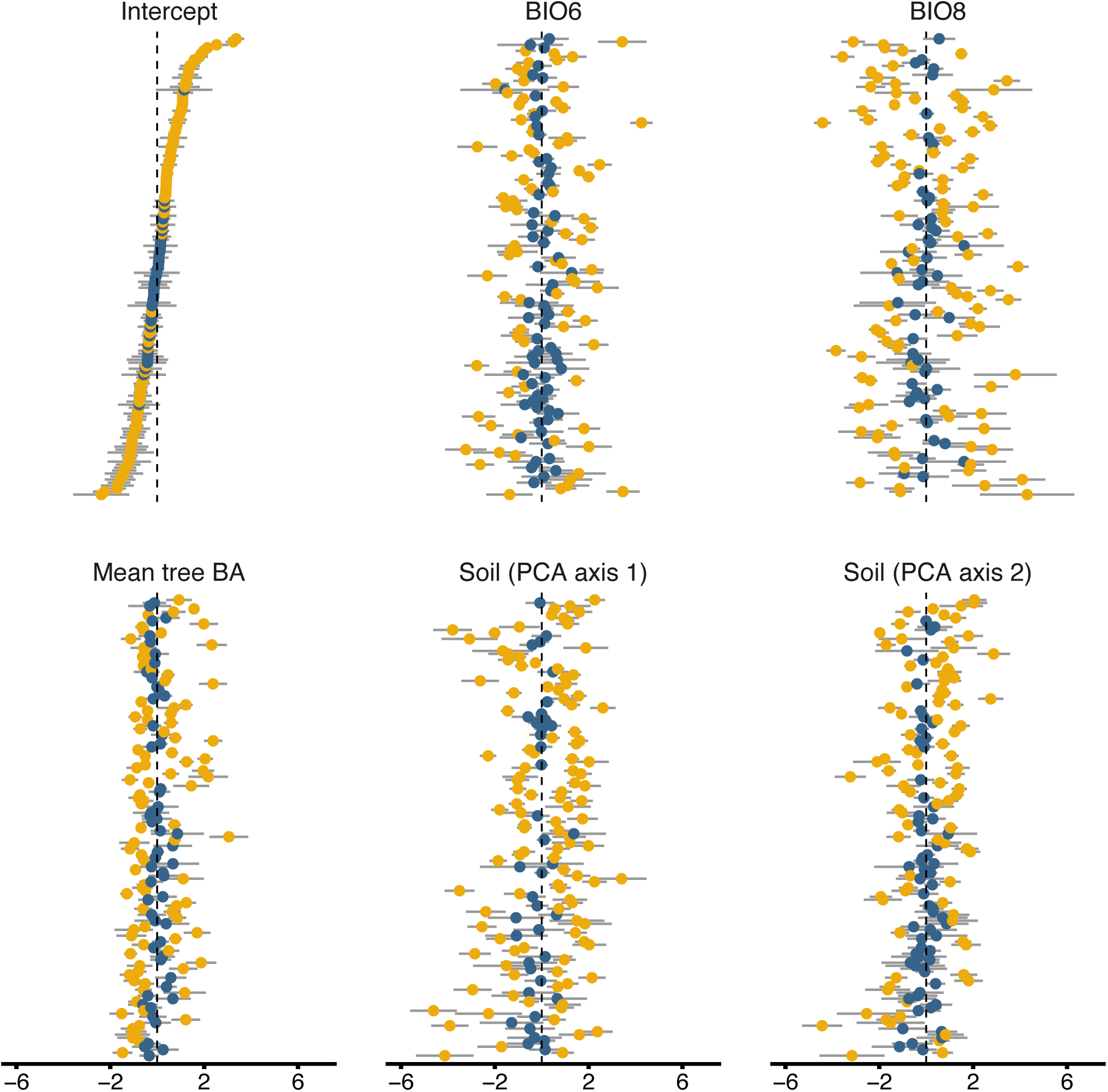
Estimated species’ abundance intercepts and environmental responses for the static 1950s model. The *Intercept* panel shows how far species depart from the mean abundance (species arranged in order of overall abundance in all panels). Other panels show how far species depart from the expected response for each of the five most important environmental predictors in the 1950s. The gray bars are 95% “confidence intervals” (based on conditional modes and variance) with yellow points indicating species for which that interval fails to overlap zero. Note the highly variable responses among species largely unrelated to overall abundance.

**Figure 4.**
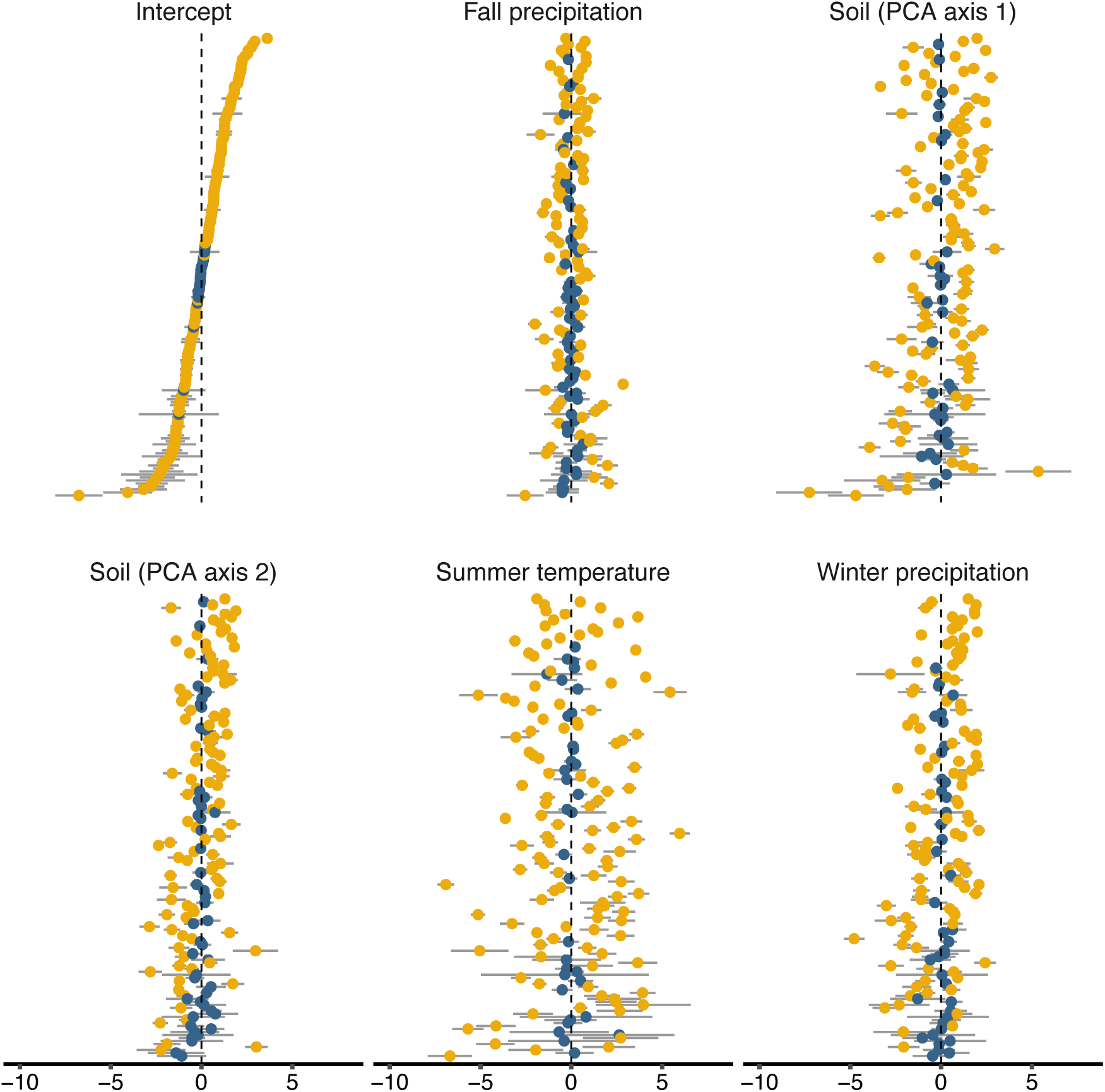
Estimated species’ abundance intercepts and environmental responses for the static 2000s model. Other details as in Fig. 3. In contrast to the 1950s model (Fig. 3, Table 3), note that less abundant species generally had more variable responses to these environmental variables.

Species’ functional traits mediate species-environment relationships judging from the plentiful trait-environment interactions evident in both the 1950s and 2000s (Fig. 5). The magnitude and nature of these effects vary greatly. Species’ area of occurrence strongly influenced their responses to climate in the 1950s (Fig. 5a). Widespread species declined more in areas with higher mean temperature in the wettest quarter (BIO8) but increased in abundance at sites with warmer minimum winter temperatures (BIO6). Widespread species tended to increase with cation concentration (Soil PCA axis 1) and finer textured soils (Soil PCA Axis 2) as expected if they are adapted to fertile sites. Species with thicker leaves declined as soil cations increased (Soil PCA1) suggesting they follow the opposite trend. Thicker-leaved species also declined under dense tree canopies (high tree BA) as expected given that thinner-leaved species are generally better adapted to shade. These species also increased with warmer minimum temperatures during the coldest month (BIO6). Species with high leaf dry matter content also tended to decline at warmer sites with richer soils.

**Figure 5.**
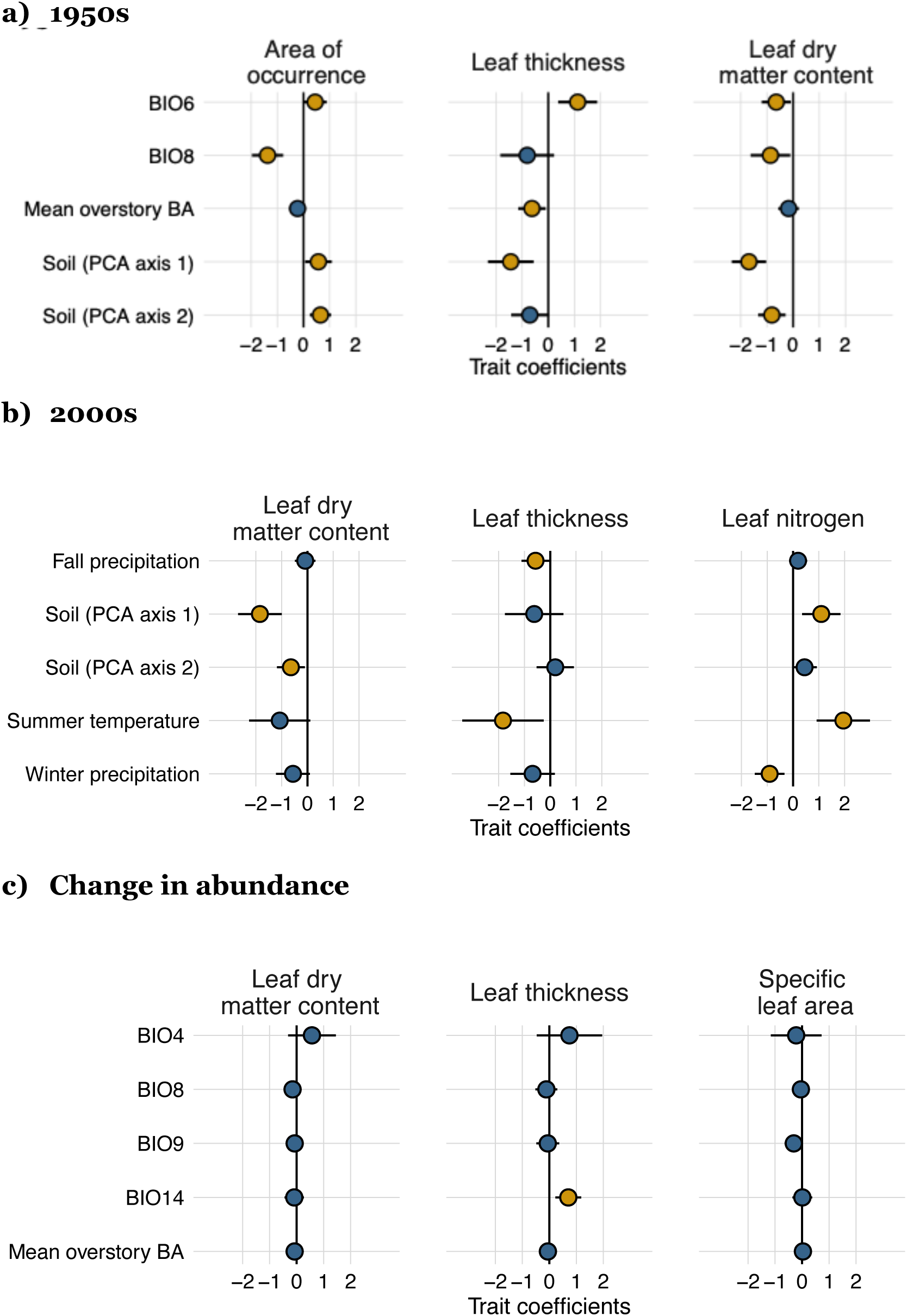
Estimated trait by environmental interactive effects in the static and dynamic models. Values show the degree to which each of the three important traits judged most important in that model modify species’ responses to significant environmental variables. (a) 195os model; (b) 2000s model, and (c) the dynamic change in abundance model. The gray bars show 95% confidence intervals with yellow points again indicating where the interval does not overlap zero.

In the 2000s, species with thicker leaves declined in areas with more fall precipitation and warmer summers (Fig. 5b). Leaf dry matter content can mediate species’ responses to soil gradients. Species with higher leaf dry matter content occurred more commonly in sandier soils (Soil PCA Axis 2) and lower cation concentrations (Soil PCA Axis 1). Species with higher leaf nitrogen tended to occur on richer soils (as expected) and higher summer temperatures but declined in areas with more winter precipitation.

### Do factors that determine abundance remain stable?

Somewhat. As already noted, many environmental variables affected species similarly in the 1950s and 2000s (Fig. 2). Mean tree basal area, however, declined in importance, probably reflecting succession toward denser canopies by the 2000s, reducing variation in this predictor. In contrast, several precipitation variables of minor significance in the 1950s became more important by the 2000s (cluster in lower left, Fig. 2). Applying the 1950s model to the 2000s data shows similar coefficients for many estimated trait-environment coefficients (Fig. 6a) including the key gradients in soil cations and texture (Soil PCA1 and PCA2). This stability suggests that soil conditions have not changed greatly at these sites since the 1950s (though our lack of soil data from the 1950s prevents us from directly testing this). Interactions of the three dominant traits with the two climate variables (minimum temperature of the coldest month and mean temperature of the wettest quarter – BIO6 and BIO8), however, showed more divergence between the 1950s and 2000s. These include area of occurrence and leaf thickness by BIO6 and BIO8 which all lost significance. In contrast, species with less leaf dry matter content tended to respond more strongly to BIO6 and the soil gradients in the 2000s.

**Figure 6.**
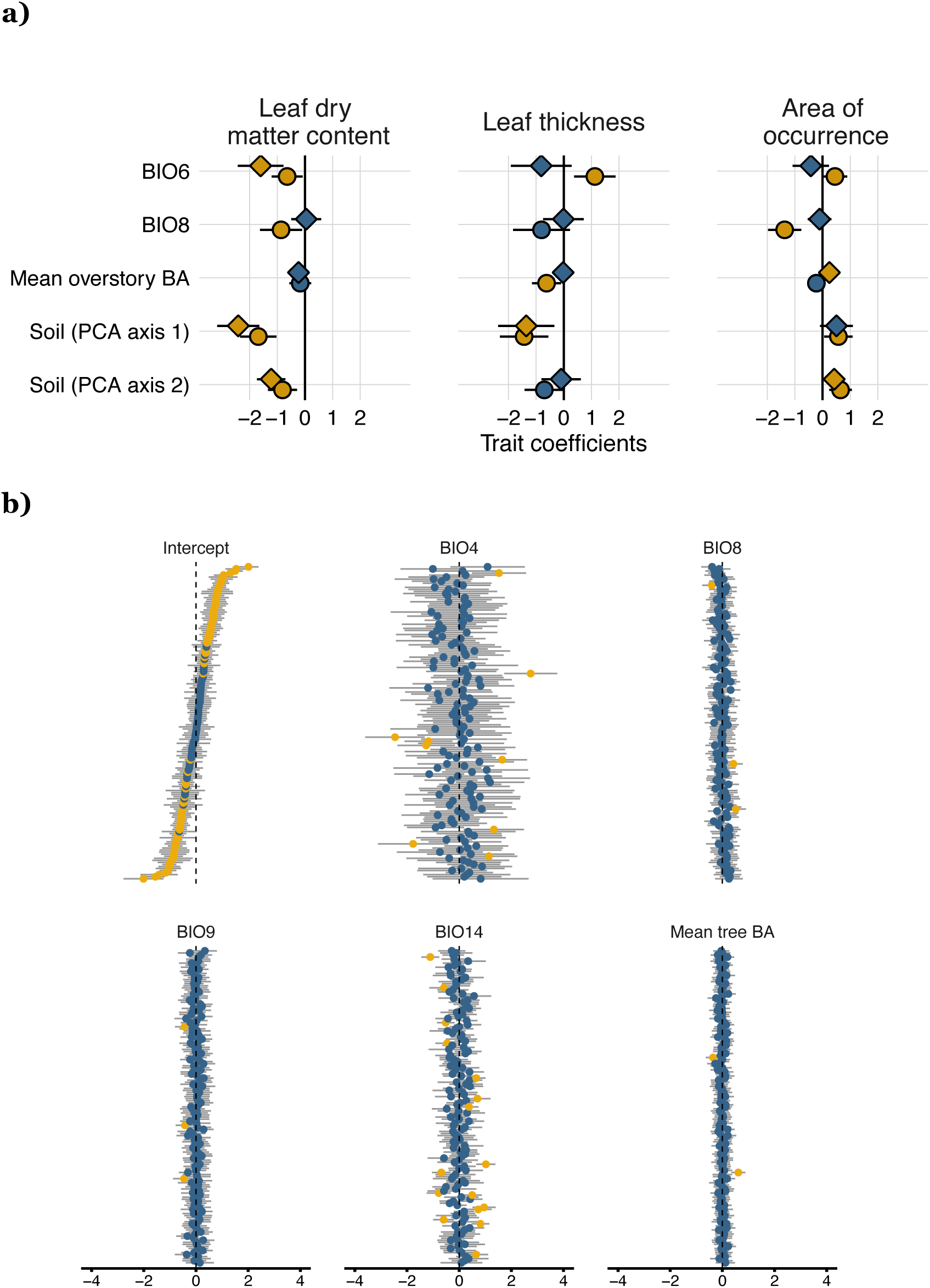
**a)** Trait coefficients estimated by applying the 195os model (circles) to the 2000s species’ abundance data (diamonds). The gray bars show 95% confidence intervals and yellow points show when that interval fails to overlap zero. Note that species respond similarly to many trait x environment effects in both the 1950s and 2000s but have started to diverge in their responses to changes in climate (BIO6 and BIO8). **b)** Results from the change in abundance model. Intercepts (upper left) show estimated mean changes in abundance across sites for each species (logit-transformed). Species are arranged in order of increasing changes in abundance (shown as departures from the grand mean). Succeeding panels show individual species departures from the overall expected response (of all species) to changes in the five most important environmental variables affecting changes in species abundance. Gray bars: 95% confidence intervals; yellow points: significantly different from zero (i.e., affected that species’ change in abundance). Changes in precipitation in the driest month (BIO14) had the most significant effects on how much species changed in abundance between the 1950s and 2000s.

### Which climatic factors affect changes in species’ abundance?

Changes in mean tree basal area at a site and shifts in many bioclimatic variables strongly affected changes in species’ abundance (Fig. S3). Changes in mean tree BA were generally positive, reflecting succession to denser canopies and shadier, moister understories in these largely undisturbed stands. As expected, these understory species were quite sensitive to these shifts in light and moisture. Neither of the two most important climate predictors for the static analyses (low temperatures in the coldest month or mean temperatures in the wettest quarter – BIO6 and BIO8) had strong effects on shifts in plant abundance. Instead, the three most important climatic factors all reflect conditions in the driest month or quarter of the year: precipitation in the driest month (BIO14), precipitation in the driest quarter (BIO17), and temperatures in the driest quarter (BIO9). This result also emerged in the hierarchical change in abundance model (Fig. 6b). Changes in precipitation in the driest month (Feb., BIO14) had more significant effects on how much species changed in abundance between the 1950s and 2000s than any other climate variable. These results suggest that species responding the most to shifting environmental conditions respond particularly to increases in winter precipitation.

### Do traits mediate species responses to changes in climate?

As already noted, models for the separate survey periods indicate that traits tend to both directly affect species’ abundances and to indirectly affect abundance by modifying how key environmental predictor variables affected abundance. These interactive effects were less evident in the combined hierarchical model analyzing changes in species abundance. In line with the simpler model discussed above, increases in precipitation in the driest month (in mid-winter, BIO14) tended to increase the abundance of species between the 1950s and 2000s (Table 5). BIO14 was the only significant climatic variable. Likewise, among traits, only leaf thickness emerged as significant, reducing the abundance of thicker-leaved species between the 1950s and 2000s. Given these limited effects, it is not surprising that the only significant trait-climate interaction (of 15 tested) was LT x BIO14. Their positive interaction implies that thicker-leaved species decreased less at sites where precipitation increased in mid-winter. Their interaction had similar magnitude and significance as the individual effects of BIO14 and LT demonstrating at least that this one trait mediates how species respond to climate changes. Other functional traits hardly affected species-climate relationships (Fig. 5c). The lack of 1950s soil data prevented us from testing how changes in Soil PCA1 and PCA2 might have affected species’ abundances.

**Table 5.**
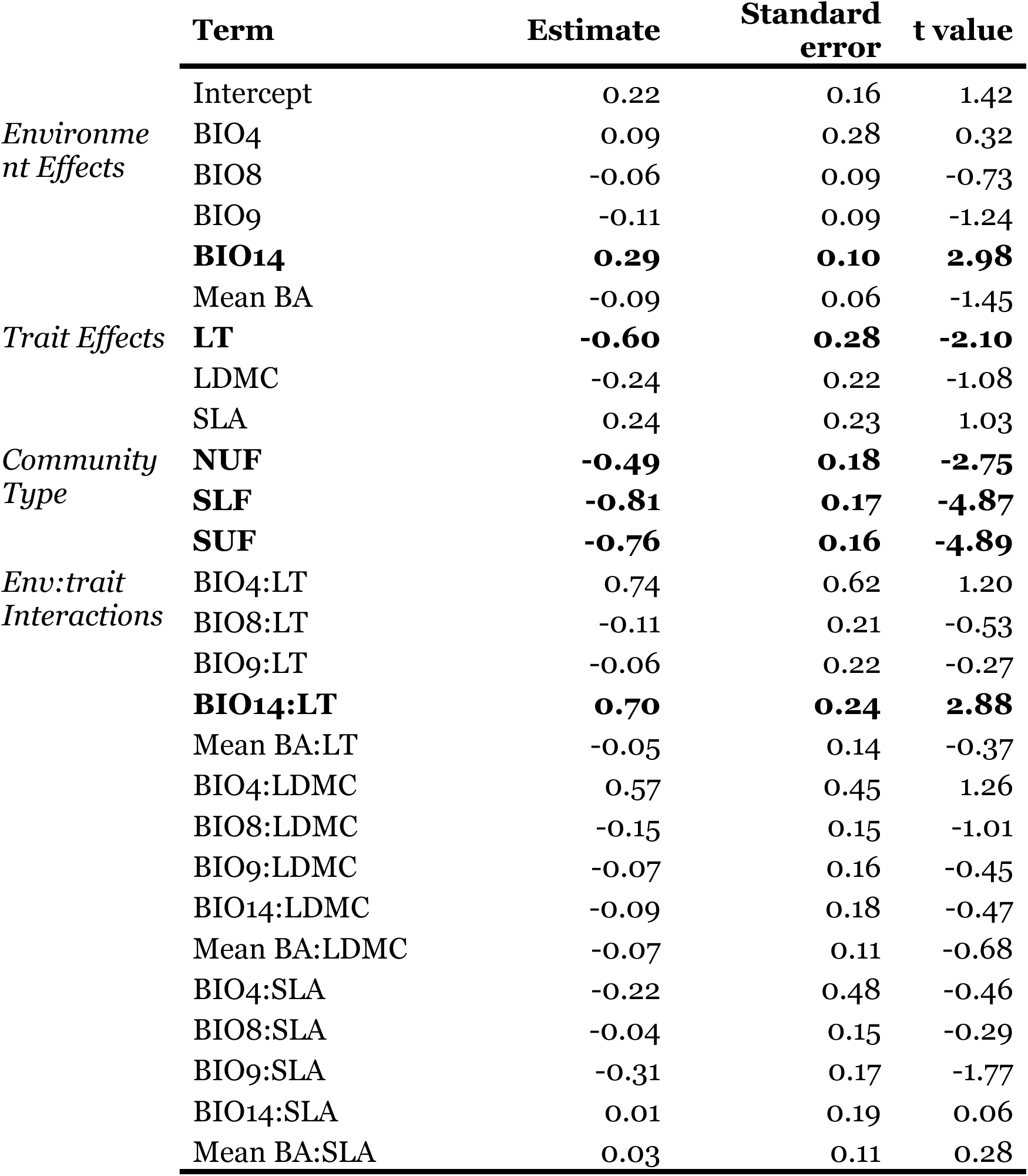
Model coefficients for the change in abundance model. Significant estimates (where confidence intervals do not overlap zero) are bolded. For environmental abbreviations, see Table 1. Trait abbreviations are: LDMC: leaf dry matter content; SLA: specific leaf area; LT: leaf thickness.

## Discussion

A classic tenant in ecology is that species respond individualistically to variation in environmental conditions reflecting the traits they evolved that adapt them to these conditions (Whittaker 1956; Curtis 1959; Maguire *et al*., 2015). Our results clearly confirm that species respond to environmental factors individually, sometimes in complex ways (Figs. 3-4) as do other studies (Ackerly *et al*., 2010; Chen *et al*., 2011; Grenouillet & Comte, 2014; Rapacciuolo *et al*., 2014). Corollaries of this tenant are that we should be able to relate variation in a species’ abundance across sites to variation in environmental conditions and variation in abundances among species to their traits. Knowing which factors affect the location and abundance of species helps explain contemporary ecological patterns and might allow us to predict how species are likely to respond to shifts in climate and other conditions in the future (Green et al. 2022). This assumes, of course, that trait-environment-abundance relationships remain stable. As the kinds of change multiple and rates of environmental change accelerate, our ability to predict these changes will likely decline. Rates of climate change are accelerate in our region (WICCI, 2011), increasing the discordance between historical and future climates (Ordonez *et al*., 2014). This may help explain why interactions between species’ traits and shifting environmental conditions are growing more complex.

We rigorously tested these assumptions and predictions using a remarkable set of data from temperate forest understories in north-central U.S. (Waller et al. 2012). These data involve careful and extensive surveys and re-surveys of the same sites over 50 years, encompassing a large number of species, sites, traits, and environmental variables (153, 284, 18, and 33, respectively). These resurveys reveal considerable variation with most species experiencing appreciable shifts in abundance between the 1950s and 2000s (see intercepts, Fig. 6b). This variation (plus state-of-the-art hierarchical statistical models correctly employing all these data) lent power to these tests. We first asked how well species’ trait values and site environmental conditions serve to predict how species vary in abundances. We confirmed that many such relationships exist, confirm those recently obtained by Rolhauser et al. (2021) and Beck et al. (2022) using similar data but distinct methods. Soil and overstory (light) conditions emerged as most important, though many climate variables also exert strong effects on species distributions. This study differs from theirs in asking whether these relationships have remained stable in these forests over the past 50 years, whether the roles of particular climate conditions and trait values have shifted in importance, and how trait-environment interactions affect these dynamics. In short, trait-environment-abundance relationships appear to be fairly stable, though the relative importance of overstory conditions may be declining and precipitation patterns appear to be growing in importance relative to temperatures. However, these relationships proved to be quite limited in terms of their utility for predicting shifts in plant abundance between the 1950s and 2000s. Why?

An earlier paper (Ash et al. 2019) demonstrated that forest plant species in Wisconsin have approximately tracked local shifts in climate except that their ability to track these changes is limited, creating moderate to large lags in most species. These lags reflect the slow life-histories of most forest species and how forests tend to ameliorate the hotter and drier conditions accompanying climate change (De Frenne et al. 2013). In addition, many other factors also affect plant populations including stochastic forces related to dispersal. These have become more important in Wisconsin forests (Rogers et al. 2008, 2009).

Our results confirm that soil and overstory conditions had strong effects on plant distributions in the 1950s and remain strong determinants of abundance in the 2000s. Plants continue to respond to climatic factors much as they responded 50+ years ago except that precipitation has become more important than temperature. Variation in overstory density (estimated crudely here by mean tree basal area) appears to affect species less in the 2000s than it did in the 1950s. This may reflect successional trends that are closing canopies in the Central Sands and southern upland forests (Rogers *et al*., 2008; Li & Waller, 2015). Large declines in mean basal area and its variation at the CSP sites (means and S.D.’s declined from 0.162 to 0.05 and 0.16 to 0.02, respectively) greatly reduced its power for predicting species’ responses.

How species respond to changes in environment is often mediated by life history characters and ecological strategies, yet these relationships can be difficult to characterize (Pollock *et al*., 2012; Jamil *et al*., 2013). Environmental conditions beyond those we characterized and shifts in landscape context alter the forces affecting plant distributions and abundance. The factors structuring species’ distributions in previous time periods may not match those affecting current distributions and those affecting shifts in distribution may also differ, as we found. Similar yet distinct combinations of traits, environmental factors, and their interactions structured species’ distributions in the 1950s and 2000s in Wisconsin forest communities. Curiously, neither set of predictors for static distributions within each period emerged as important for predicting changes in plant abundance over time (see below).

### Environmental factors and traits affect species distributions

Environmental gradients strongly affected the distributions and abundances of these forest species and functional traits mediated species responses to these gradients. Similar sets of variables reflecting soil and overstory conditions, growing season temperatures, and the water/energy variables influenced species’ distributions in both the 1950s and the 2000s (Fig. 2, Tables 3-4). However, temperature and precipitation patterns shifted considerably across the state over the past 50+ years (Kucharik *et al*., 2010). This may help explain why precipitation-related variables increased in importance by the 2000s. Simultaneously, links between understory communities and overstory dynamics have weakened in southern forests while landscape factors including habitat fragmentation have grown stronger (Rogers *et al*., 2009). Changes in land cover and use precede the original Curtis surveys with forests cut over in the North between 1870 and 1920 continuing to succeed to denser conditions and more open oak-hickory forests in southern Wisconsin continuing to grow denser canopies (Rhemtulla *et al*., 2009). These changes help to explain the strong successional signal we found and the declining importance of tree basal area by the 2000s.

Despite pronounced changes in overstory and climatic conditions, these failed to accurately predict changes in species abundances at these sites. As noted above, these species have already shifted their distributions in response to changes in local climatic conditions across Wisconsin with species’ centroids tracking these changes but usually lagging them (Ash *et al., 2017*). These lags probably reflect the slow life histories of many forest plants, the ability of forest canopies to ameliorate the effects of climate change (De Frenne *et al*., 2013), and other factors affecting species dynamics including herbivory (especially by deer in our region), habitat fragmentation, invasions by weedy plants and exotic earthworms, and nitrogen deposition. Without data on these factors, it is impossible to compare their separate and combined effects to those of climate change.

### How do functional traits mediate species-environment relationships?

We identified several key traits mediating effects of environmental gradients on species’ abundances as did Pollock et al. (2012) and Jamil et al. (2013) using similar approaches. These results extend those from an earlier study that also found persistent trait-environment relationships in the upland subset of the sites we examined (Amatangelo *et al*., 2014). Our more sophisticated statistical approach allowed us to analyze how species’ abundances depend not only on individual traits and environmental factors but also how traits modulate species’ responses to environmental gradients. Future work in other systems confirming these relationships would broaden our understanding of which combinations of traits and environmental factors reliably serve to predict variation in species abundance.

The area of occurrence and leaf traits strongly affected species-environment relationships in the 1950s (Fig. 5a). This is not surprising given that widespread, habitat generalists tolerate a wide set of environmental conditions (Warren *et al*., 2001). Species with narrow distributions often have more specialized habitat requirements (***cites***). Both characteristics make specialists more vulnerable to nitrogen deposition (Staude et al. 2020). Similarly, we found more broadly distributed plant species to be more efficient at tracking changes in climate than narrowly adapted, more local species (Ash *et al., 2017*).

Species with higher leaf dry matter content occurred more commonly in coarse, low fertility soils and at sites with colder winters (BIO6) and summers (BIO8). Such species also tend to have longer leaf lifespans, higher investment in structural defenses and support, and more conservative physiological and life history characteristics (Givnish, 1979; Cornelissen *et al*., 2003). Declines in thicker-leaved species in forests with denser overstories probably reflect shadier conditions at these sites (Rogers *et al*., 2008; Johnson *et al*., 2014; Li & Waller, 2015). These thick-leaved species tend to have high leaf matter content and to occur in low fertility soils (Pérez-Harguindeguy *et al*., 2013). These traits confer advantages in colder climates and in sites with lower fertility, common in northern Wisconsin. Oddly, in contrast to species with high leaf dry matter content, thicker-leaved species declined with colder winter temperatures, highlighting the complexity of trait-environment interactions.

Species with thin, high nitrogen leaves occurred more abundantly in areas with higher summer temperatures, lower winter precipitation, and higher soil fertility. Such conditions are common in southern Wisconsin supporting high photosynthetic rates. Summer and winter temperatures and winter precipitation are all projected to increase in Wisconsin with the ratio of snowfall to rain declining (Notaro *et al*., 2010). How these changes will affect plant communities remains unclear. Few studies have examined the ecological impacts of changing winter climates in temperate regions (Kreyling, 2010; Ladwig *et al*., 2016). Nonetheless, our models suggest that species with high SLA and leaf N in southern Wisconsin are poised to increase greatly in response to current shifts in climate (and possibly N deposition – Simkin et al. 2016). However, forest sites in southern Wisconsin have also experienced considerable habitat loss and fragmentation plus widespread fire suppression, increasing rates of local extinction (Rogers *et al*., 2009). Climatic changes interact with these other changes in conditions to substantially alter the species and functional composition of communities (Li & Waller, 2017).

### Have trait-environment relationships shifted?

Our 1950s model served adequately to predict 2000s abundances (Fig. 6a). This suggests consistent trait-mediated responses to environmental gradients. A few basic traits had important effects modulating how species respond to shifts in soils, the canopy, and climate. This suggests optimism regarding our ability to predict species’ responses to future environmental change. However, how traits interacted with January minimum temperatures (BIO6) and temperatures during the wettest quarter (BIO8, usually summer) shifted, losing significance by the 2000s. Seasonal temperatures have changed dramatically since the 1950s with winter months warming considerably while summer temperatures have actually decreased in parts of the state (Kucharik *et al*., 2010). These large changes could disrupt underlying species-environment relationships (Fig. S4), reducing our ability to predict changes in species abundance (over and above the lags we observe in the ability of many species to track these shifts in climate – Ash *et al.,* 2017).

Although other studies found that functional traits serve to predict species-climate relationships (Soudzilovskaia *et al*., 2013; Li *et al*., 2015), our trait-environment model had limited power for predicting long-term changes in species’ abundances (Fig. 4c, Fig 6b). Leaf thickness alone modified how species responded to mid-winter precipitation. This may imply that other, unmeasured traits or local or landscape environmental factors had stronger effects on shifts in species abundances. Simultaneous changes in multiple environmental factors, occurring at different rates and scales, further limits our ability to isolate and identify key factors affecting plant population/meta-population dynamics. These simultaneous changes have likely created novel combinations of conditions never experienced by these species (Radeloff et al. 2015). In addition, as stochastic forces and local dispersal rise in importance (as documented in SUF), species-environment relationships surely weaken. These complications reduce our ability to predict changes in plant distribution and abundance but this should not deter us from seeking to improve our ability to predict species’ responses to the drivers of ecological change.

## Acknowledgments

This project was supported by the following National Science Foundation grants: DEB 9974041, DEB 0236333, and DEB-0717315, and a National Research Initiative grant #2008-35320-18680 from the USDA National Institute of Food and Agriculture Biology of Weedy and Invasive Species Program. A grant from NSF’s Dimensions of Biodiversity program (DEB-1046355) supported collection of the trait data and resurveys of the Pine Barrens sites.

## Appendix

**Figure S1.**
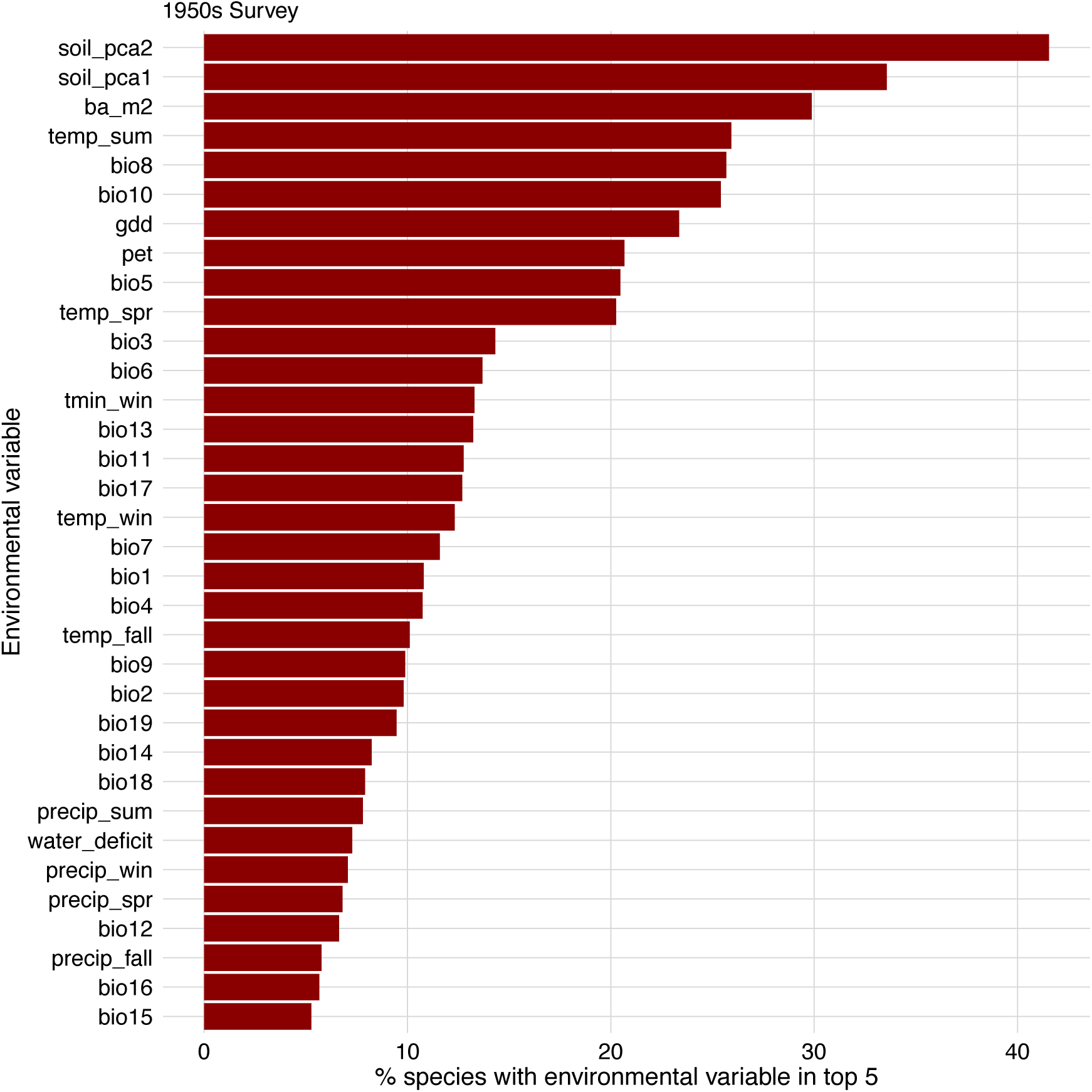
The relative importance of all 34 environmental variables for explaining the distribution of species in the 1950s survey based on random forest models. Random forest models were generated for each species and environmental variables are ranked according to the percentage of models where the variable was one of the top five most important variables.

**Figure S2.**
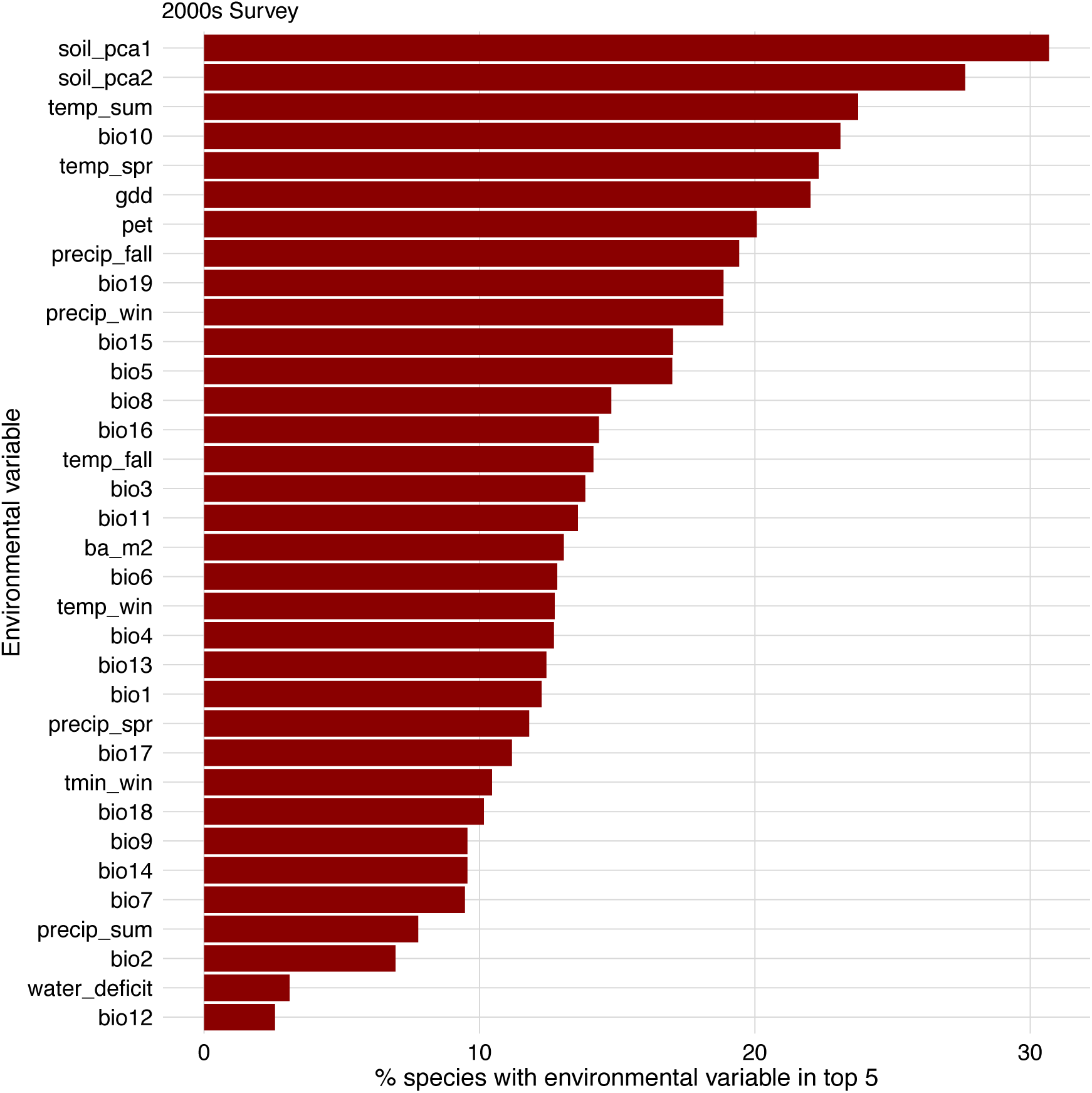
The relative value of all 34 environmental variables for predicting the abundances of species in the 2000s survey as inferred from the random forest models. Models for each species analyzed how well these variables predicted abundances in the early 2000s. Environment variables were then ranked according to the percentage of models that included that variable among the five best predictors. For abbreviations, see Table 1.

**Figure S3.**
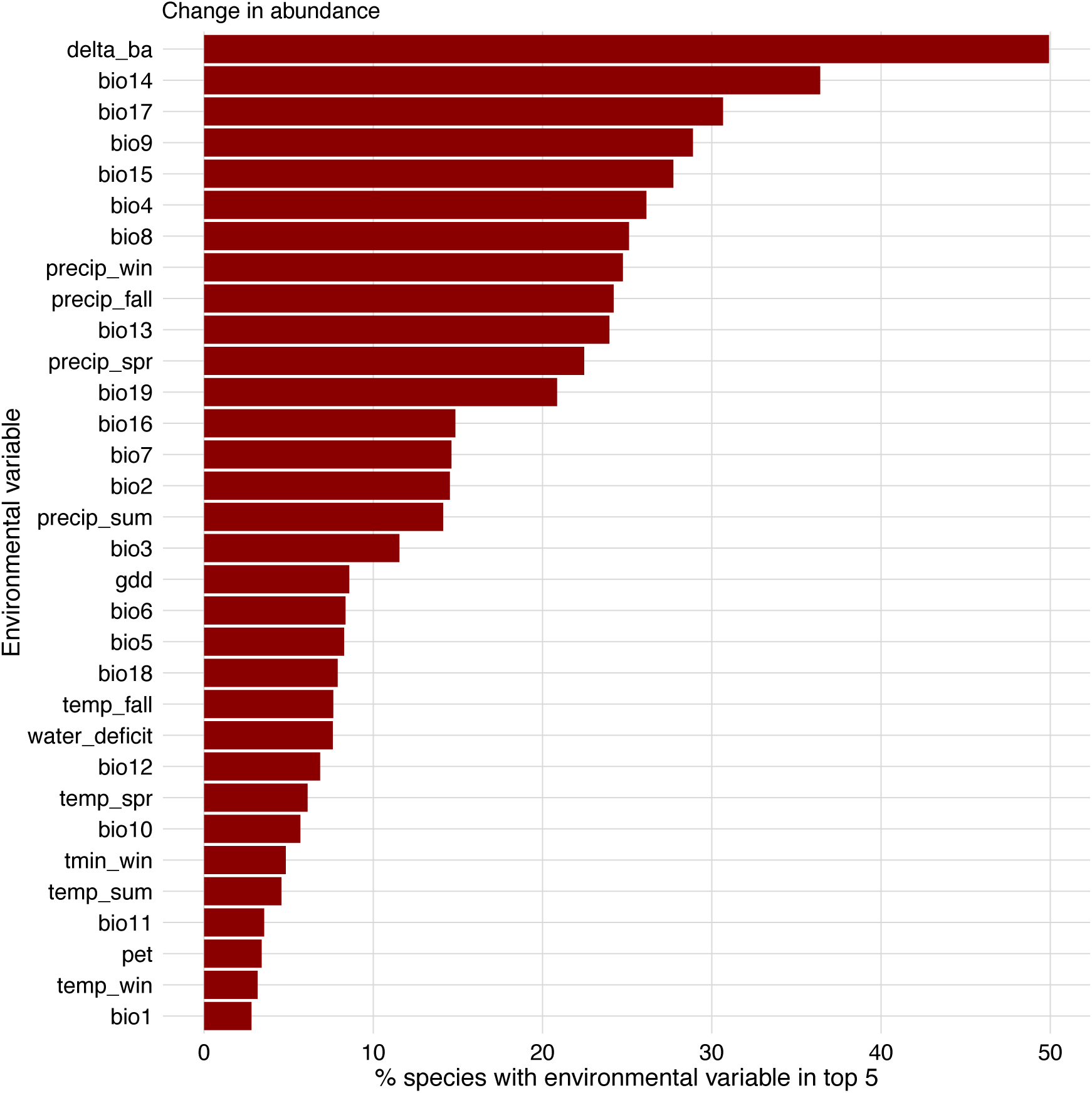
The relative importance of changes in mean tree basal area (delta_ba) and 31 bioclimatic factors for predicting changes in the abundance of species as inferred from the random forest models. Models for each species analyzed how these environmental variables affected 1950s-2000s changes in abundance. Note that soil variables were excluded from these analyses. Environment variables are ranked according to the percentage of models that include that variable within the five best predictors. For abbreviations, see Table 1.

***<< Given that this is a primary focus of the paper, should we promote this Fig. to the main text? >>***

**Figure S4.**
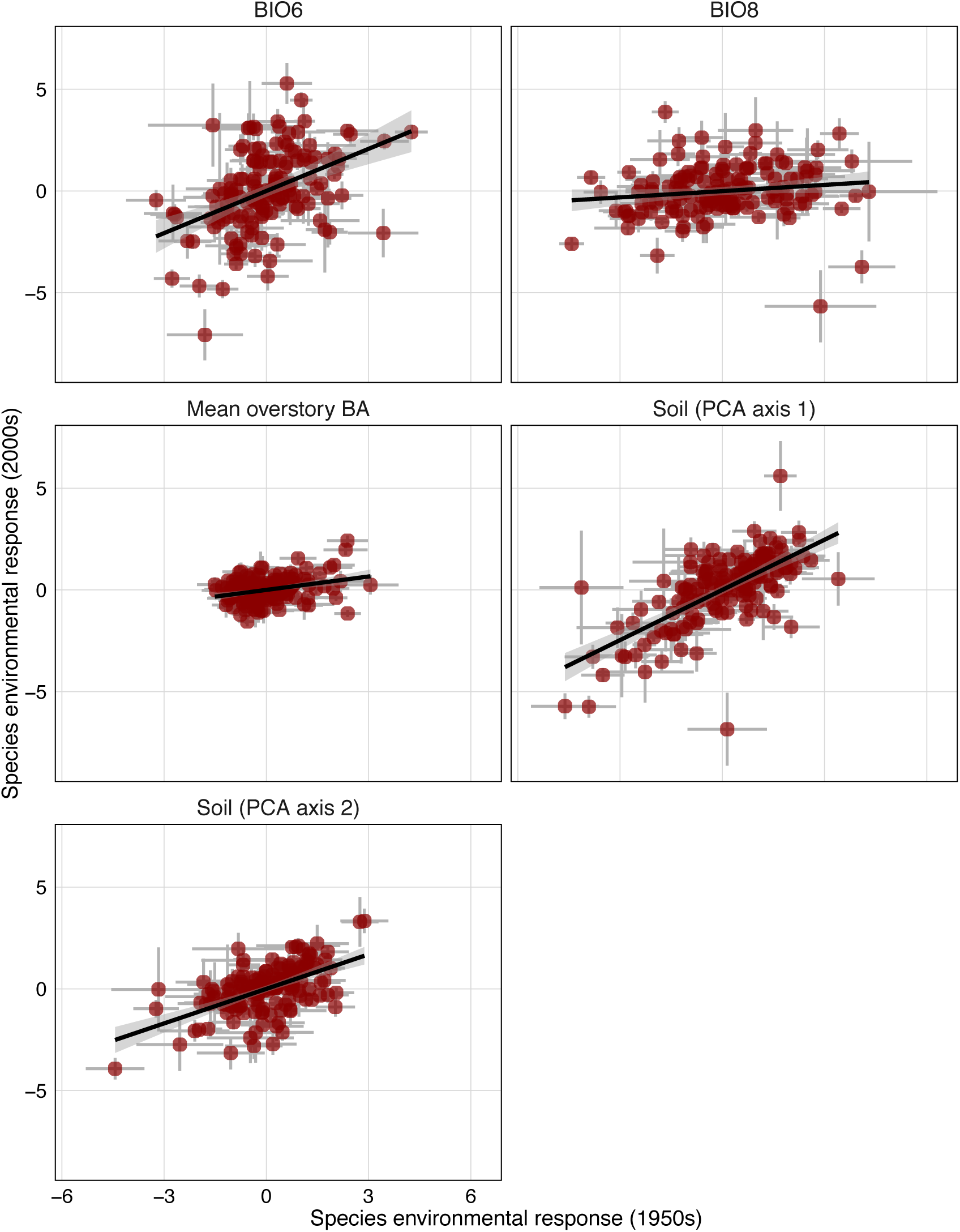
The relationship between the estimated species-environment response intercepts for the 1950s model (as in Fig. 3a) and when the 1950s model structure is applied to the 2000s data. Relationships are shown for each for the environmental predictor variables. The gray bars are 95% confidence intervals. Note the increased variation for the BIO6 and BIO8 relationships. These two variables have experienced larger changes over time, relative to other climate change trends.

